# Engineered Riboswitch Nano-carriers as a Possible Disease-Modifying Treatment for Metabolic Disorders

**DOI:** 10.1101/2022.05.16.492066

**Authors:** Shai Zilberzwige-Tal, Danielle Gazit, Hanaa Adsi, Myra Gartner, Rahat Behl, Dana Laor Bar-Yosef, Ehud Gazit

## Abstract

Both DNA- and RNA-based nanotechnologies are remarkably useful for *in vitro* molecular-scale device engineering and are applied in a vast array of applications. However, while the function of nucleic acid nanostructures is robust under *in vitro* settings, their implementation in real-world conditions requires overcoming their inherent degradation sensitivity and subsequent loss of function. Viruses are minimalistic yet sophisticated supramolecular assemblies, able to protect their nucleic acid content in inhospitable biological environments. Inspired by this natural ability, we engineered RNA-virus-like particles (VLPs) nanocarriers (NCs). We showed that the VLPs can serve as an excellent protective shell against nuclease-mediated degradation. We then harnessed biological recognition elements and demonstrated how engineered riboswitch NCs can act as a possible disease-modifying treatment for genetic metabolic disorders. The functional riboswitch is capable of selectively and specifically binding metabolites and preventing their self-assembly process and its downstream effects. When applying the riboswitch nano-carriers to an *in vivo* yeast model of adenine accumulation and self-assembly, significant inhibition of the sensitivity to adenine feeding was observed. In addition, using an amyloid-specific dye, we proved the riboswitch nano-carriers ability to reduce the level of intracellular amyloid-like cytotoxic structures. The potential of this RNA therapeutic technology does not stop at metabolic disorders, as it can be easily fine-tuned to be applied to other conditions and diseases.

---

The straightforward Watson-Crick base pairing has enabled exceptional control over the shape, size, and geometry of both DNA and RNA nanostructures.^1–4^ This, has led to the rapid development of the nucleic-acids nanotechnology field with numerous applications.^3,5–10^ Yet, one of the greatest limitations of nucleic acid-based devices is their rapid nuclease-mediated degradation, when applied *in vivo*.^4,11^ One way to overcome this hurdle is to chemically modify the nucleic acids which are costly. Additional way is to encapsulate the nucleic acids in liposomes or lipid nanoparticles.^12–15^ However, in some cases it is necessary to retain the nucleic acids structures’ functionality for specific applications and to enable passive diffusion of molecules inside the particles. Both liposomes and LNPs do not pose these features.

Riboswitches are RNA-based genetic control elements that are usually found in non-coding regions of RNAs.^16,17^ The riboswitches are highly structured and able to bind target cellular metabolites with high specificity.^18^ Yet, similar to other functional RNA applications, this technology is significantly limited by the high sensitivity of the RNA molecules to the host nucleases.^4^ Aiming to address this issue, several approaches have been taken, including liposomes and LNPs, mostly utilizing various protective coatings of the RNA molecule. A prominent example that utilizes the wild-type (WT) MS2 bacteriophage is a 27 nm particle consisting of a single copy of the maturation protein and 180 copies of the coat proteins (CP) (organized as 90 homodimers) arranged into an icosahedral shell.^19–21^ The bacteriophage’s assembly is driven by a specific 20 nucleotides RNA sequence forming a stem-loop structure called “translational repression RNA” (TR-RNA), encoded in the MS2 genome.^22,23^ The final fully formed bacteriophage capsid generally represents the lowest free energy state of the component proteins and nucleic acids, leading to a stable structure.^24^

Here, we aimed to develop a disease-modifying treatment that disrupts the early stages of the metabolite self-assembly process, which is linked to metabolism disorders. For this purpose, we encapsulated a functional riboswitch inside MS2 bacteriophage virus-like particles (VLPs) (Figure 1, Figure S1).^25^ Utilizing the VLP as the nano-carrier (NC) of the riboswitch was found to provide three important features: (i) retaining the structure and therefore the function of the riboswitch, (ii) the VLPs pores permitted the passive influx of small molecules, such as metabolites, and (iii) the VLPs shell acted as a protective cage against nucleases, thus overcoming one of the most significant limitations of functional RNA applications. Furthermore, MS2 VLPs hold great potential in applying them as vaccines and as a carrier to specific cells.^26,27^

**Figure 1.**
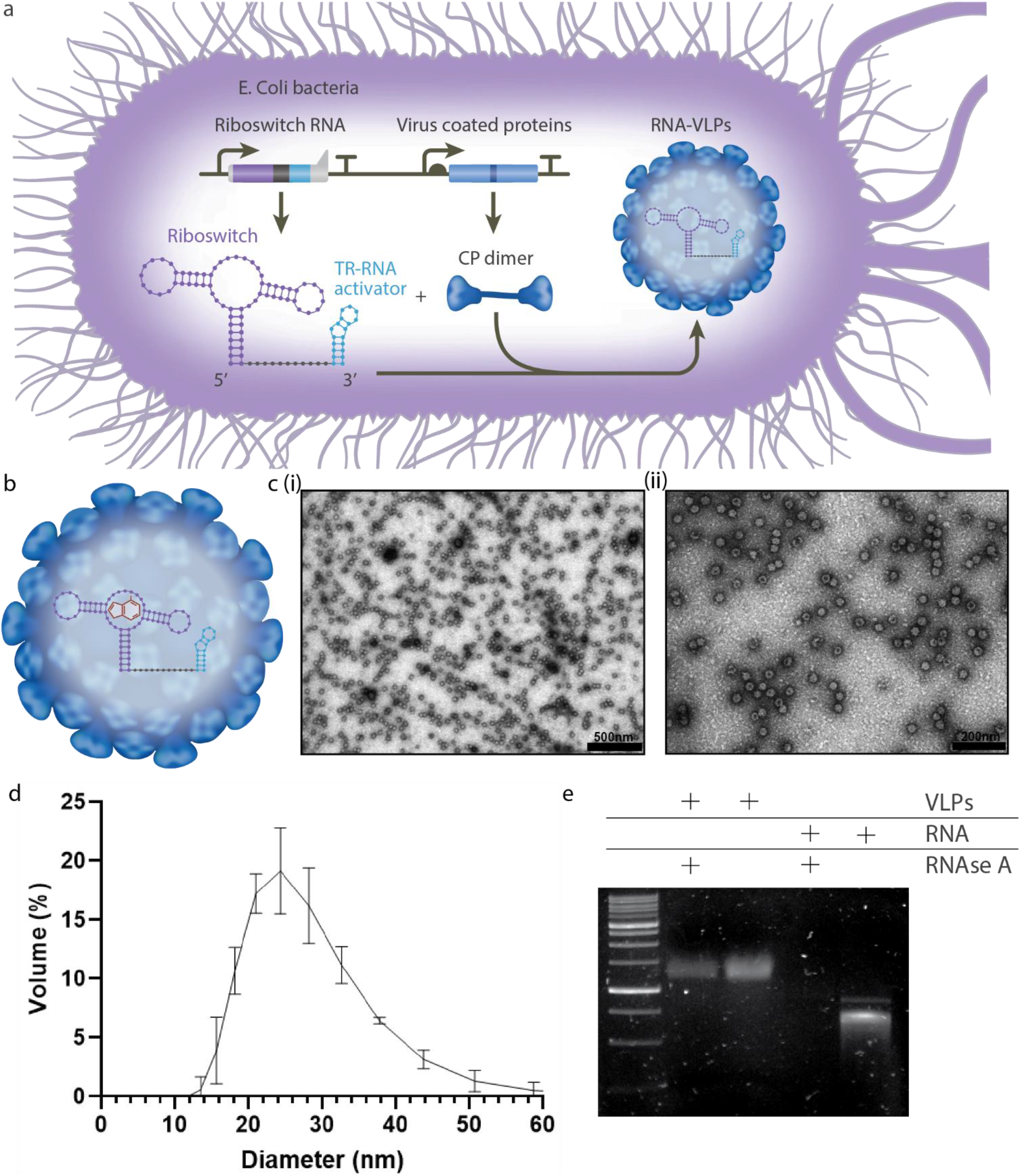
VLPs production system and characterization. a. The *in vivo* VLPs production mechanism. The system consists of: (i) A gene encoding the riboswitch RNA, and (ii) A gene encoding the virus CP, which assemble into CP dimers. The riboswitch RNA gene is designed to include at its 3’ end the TR-RNA activator sequence, leading to the VLPs’ self-assembly and the riboswitch encapsulation within the VLPs. Following their self-assembly, the Riboswitch-VLPs are purified from the bacteria. b. The Riboswitch-VLPs maintain its functionality while encapsulate and capable to bind metabolite that are passively diffused in and out of the particle. c. TEM images of the *in vivo* produced VLPs-RNA nano-carriers. Scale bar: 500nm d. Dynamic light scattering measurements of the VLPs-Riboswitch nano-carriers e. Agarose gel electrophoresis of VLPs and RNA with or without RNase A treatment for 1 hr at 37 °C.

Error of metabolism disorders arise from an inability of the cellular machinery to perform critical biochemical reactions along a biosynthetic pathway.^28^ In the case of inborn error of metabolism (IEM) disorders, most disorders are due to a genetic mutation leading to the malfunction of an enzyme crucial for a metabolic process.^29,30^ This, in turn, results in metabolite accumulation and tissue damage that can lead, by a poorly understood mechanism, to mental retardation, epilepsy, and organ damage.^31^ To date, over 1,000 IEM disorders have been reported and their relative rarity in the general population results in a limited amount of targeted research. However, jointly, IEM disorders constitute a very substantial part of pediatric genetic diseases.^32,33^ Since the molecular basis of tissue damage is mainly unknown, it leaves the patients without disease-modifying treatment.

A plethora of degenerative disorders, including Alzheimer’s disease, Parkinson’s disease, and type II diabetes, are all associated with the formation of well-ordered, self-assembled amyloid fibrils.^34–36^ The pioneering work by Dobson and co-workers revealed that proteins not related to disease, such as myoglobin, could also form typical amyloid structures.^37,38^ Moreover, induction of apoptotic cell death by amyloid assemblies appears to be a generic property of the formed structures rather than a disease-specific pathology, implying a common mechanism for the toxic effect.^39^

Our group recently showed that the original amyloid hypothesis could be extended to non-proteinaceous assemblies formed by small metabolites, such as single amino acids and nucleobases, which also induce apoptotic cell death.^31–33^ Furthermore, the formation of amyloid-like assemblies by metabolites associated with metabolic disorders e.g., adenine, tyrosine, and phenylalanine, and the presence of antibodies against these assemblies in patients, implies a general phenomenon of amyloid formation and offers a new paradigm for metabolic disorders.^31,32,40^ Thus, metabolite amyloids may play a key role in the development of metabolic disorders, and further research addressing metabolite amyloids may shed light on the mechanisms underlying metabolic disorders, potentially leading to possible new treatments.

In humans, adenine salvage is catalyzed by two enzymes: adenine phosphoribosyltransferase (APRT) and adenosine deaminase (ADA). Mutations in the genes encoding these enzymes can lead to the accumulation of adenine and its derivatives.^41,42^ In budding yeast, APT1 and AAH1, the APRT and ADA orthologs, respectively, are similarly involved in the adenine salvage pathway.^42^ In our yeast model, designated aahlΔ aptlΔ, both AAH1, and APT1 were disrupted. This yeast model exhibits adenine accumulation thereby recapitulating adenine IEM disorders. Using this model, we were able to show that the accumulated adenine inside the cells form amyloid-like structures recognize by specific antibodies and with the ProteoStat amyloid specific dye. Furthermore, the behavior of the cells resulted in a slower growth rate on media containing normal concentrations of adenine in comparison to media without adenine.^43,44^

We used a double-expression plasmid encoding for (i) MS2 CP dimer linked via his tag and (ii) non-coding RNA sequence composed of the TR-RNA activator sequence in its 3’ conjugated to functional RNA sequence is its 5’ (Figure 1a, Figure S1), thus eliminating the potential encapsulation of non-desired RNA intermediates. Furthremore, due to the maintained functionality of the RNA and the passive diffusion of small molecules in and out of the VLPs, the RNA can selectively bind metabolites (Figure 1b).

Co-expression of the MS2 CP dimer with the non-coding DNA containing the TR-RNA activator sequence resulted in the self-assembly of the RNA-VLPs within the bacteria. Following expression, the RNA-VLPs were purified from the bacteria (see methods) and the samples were analyzed using transmission electron microscopy (TEM) where RNA-VLPs ~30nm in diameter (similar to WT MS2) were observed (Figure1 c(i) and c(ii), Figure S2 and S3). In addition, dynamic light scattering (DLS) showed a particles population of ~27 nm (Figure1d).

The stability of the RNA-VLPs in the presence of RNase A was tested. As a control, *in vitro* transcribed naked RNA was also incubated under the same conditions. The ability of the VLPs to protect the encapsulated RNA from nuclease-mediated degradation was determined using agarose gel electrophoresis (Figure 1e). While the RNA-VLPs were resistant under these conditions, the naked RNA degraded. It should be noted that during the purification process, to eliminate endogenous *E. coli* RNA and DNA, RNase and DNase were added in access to the lysis buffer, indicating that the RNA-VLPs are also resilient to DNase. TEM analysis also allowed to verify that the VLPs were stable at 4°C for at least nine months post-production (Figure 2).

**Figure 2.**
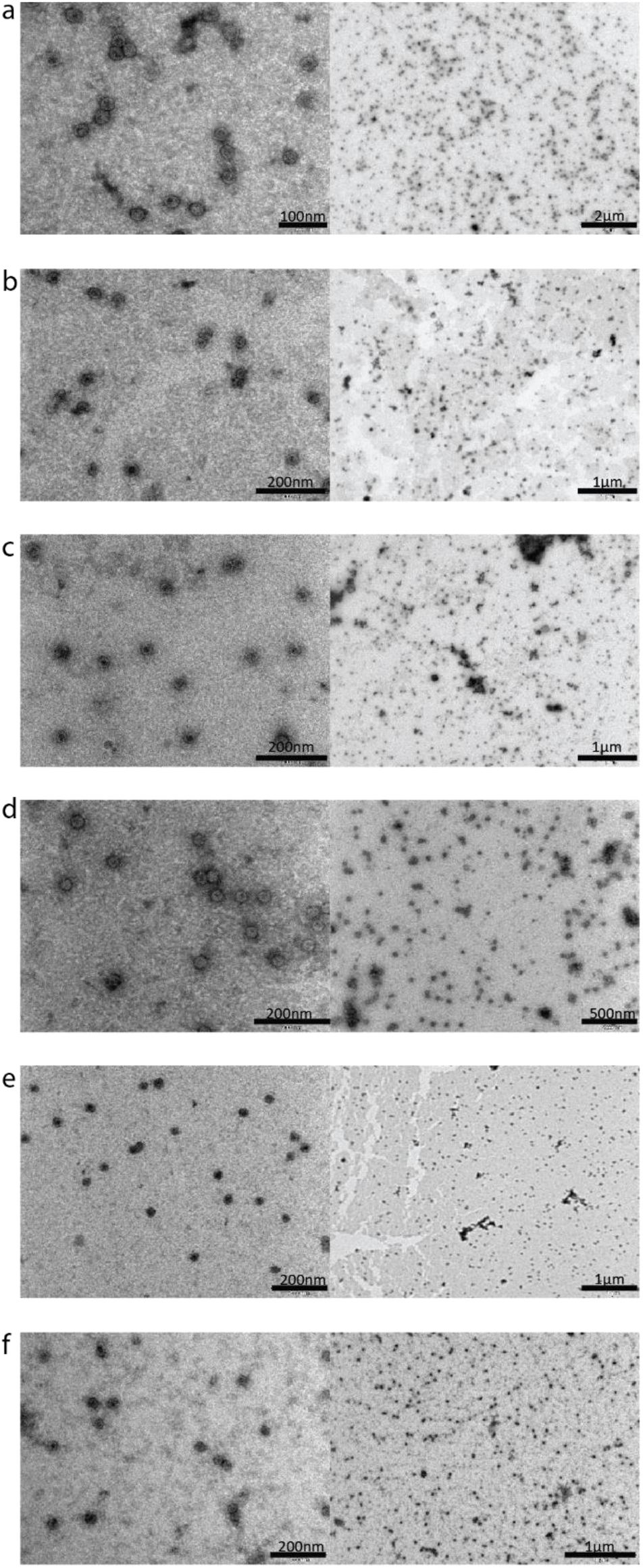
TEM images of the Riboswitch-VLPs nano-carriers at different post-production time points: a. two weeks. b. one month. c. two months. d. three months. e. six months and f. nine months.

Next, we aimed to apply the RNA-VLPs as a possible disease-modifying treatment for metabolic disorders associated with elevated concentrations of metabolites. We engineered the RNA-VLPs to encapsulate a riboswitch that can specifically and selectively bind adenine. We hypothesize that supplementing media containing adenine with the Riboswitch-VLPs will result in a rescued phenotype of the aahlΔ aptlΔ strain since the Riboswitch-VLPs will bind the adenine and prevent it from accumulating and self-assemble inside the yeast cells (Figure 3).

**Figure 3.**
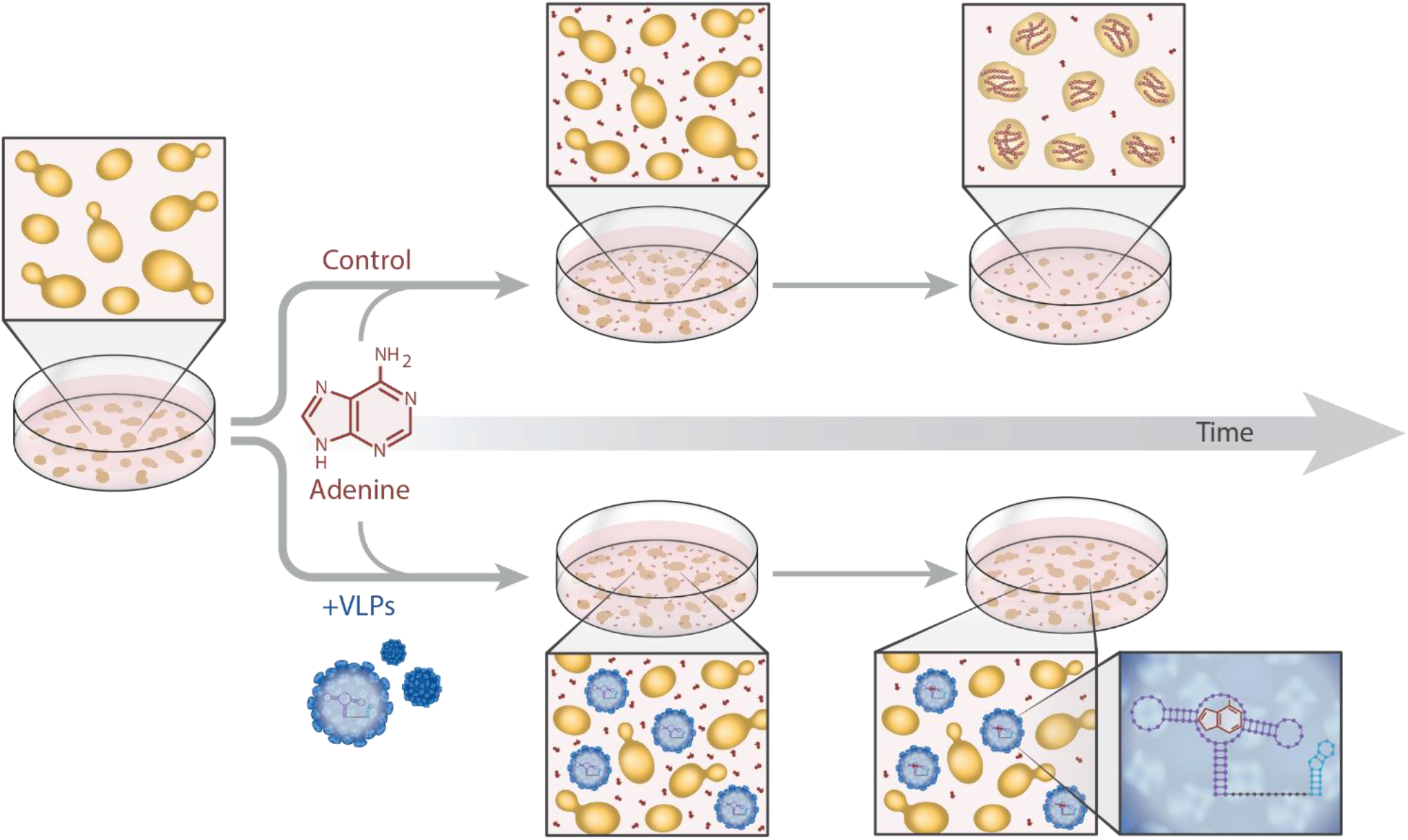
Illustration of the Riboswitch-VLPs mechanism of action. The yeast medium contains adenine and is either supplemented with the Riboswitch-VLPs or not. In the un-supplemented medium, the adenine is free to enter the yeasts where it accumulates and leads to growth inhibition. When supplemented, the riboswitch inside the VLPs can bind the free adenine in the medium, preventing it from entering and accumulating inside the yeast cells.

The natural adenine riboswitch can selectively bind adenine with Kd=354±17 nM.^18^ We chose to encapsulate a modified riboswitch with a point mutation (A58U) that increases the Kd to 273±17 nM (see supplementary information for full sequences and structures).^18^ The modified riboswitch was conjugated to the MS2 TR-RNA activator sequence and Ri-boswitch-VLPs were produced from *E.coli* as described above (Figure 1a).

The aahlΔ aptlΔ yeast model was grown in media with increasing concentrations of adenine (0, 1, 2, and 4 mg/L) and the media was supplemented with the Riboswitch-VLPs from T=0. Our results indicate that the addition of the Riboswitch-VLPs to the media significantly improved the aahlΔ aptlΔ growth in comparison to the growth without the VLPs (Figure 3a-c). The addition of the Riboswitch-VLPs alone did not affect the growth of the aahlΔ aptlΔ (Figure 3d). When the media was supplemented with VLPs encapsulating a random RNA sequence instead of the riboswitch sequence, the VLPs had no rescue effect on the yeast growth, indicating that the rescued phenotype of the yeasts resulted from the ability of the encapsulated riboswitch to bind adenine rather than the presence of the entire RNA-VLPs moieties in the medium (Figure S4). Next, we compared the growth of the aahlΔ aptlΔ mutant cells at different time points (15, and 25 hours) (Figure 3e-g). The OD600 of aahlΔ aptlΔ was significantly higher when supplemented with the Riboswitch-VLPs after 15 hours and 25 hours, at all adenine concentrations. These results further demonstrate the rescue effect of the Riboswitch-VLPs.

To quantify the rescue effect, we quantified the percentage of growth in the presence of the Riboswitch-VLPs supplementation compared to the absence of supplementation (Figure 3h). Across all adenine concentrations, there was a significant increase in the growth of the supplemented aahlΔ aptlΔ relative to the non-supplemented control. Moreover, the highest growth fold change of ~3 was measured at 4 mg/L adenine. We further tested the effect of supplementing the media with the Riboswitch-VLPs at different time points (different OD600). The Riboswitch-VLPs were found to be most effective when added at earlier time points of the yeast growth, while a less significant effect was observed when the VLPs were added at later stages (Figure S5). This may suggest that the Riboswitch-VLPs are most effective before nucleation and early oligomerization of the metabolite inside the cells, which further validates our hypothesis that the Riboswitch-VLPs can bind the adenine and prevent its intake by the yeast.

Next, we compared the effect of Riboswitch-VLPs to tannic acid (TA). TA is a widely studied polyphenol with *in vitro* anti-amyloidogenic activity against β-amyloid.^45^ We previously showed the inhibitory effect of TA on adenine both *in vitro* and *in vivo*, using the same yeast model (aah1Δ apt1Δ).^43^ In the *in vivo* model, we showed that the addition of TA to the media results in a decrease of amyloid-like structures of adenine inside the yeast cells. We previously showed that the TA can enter the cell and inhibit the self-assembly process of the accumulated adenine inside the cells.35 Similar to the assay described above, the model yeast aahlΔ aptlΔ was grown under different conditions: (i) without adenine, (ii) with adenine, (iii) with adenine, and Riboswitch-VLPs, and (iv) with adenine and TA (Figure 4). Both in the low and high adenine concentrations in the media (1mg/L and 4mg/L, respectively), the Riboswitch-VLPs and TA showed improved growth of the model yeast while in the high concentration of adenine, the Riboswitch-VLPs improved the growth also as compared to TA (Figure 4d). This result suggests that early intervention in the accumulation and self-assembly process may lead to a pronounced phenotypic rescue. TA enters the yeast and inhibits the intracellular self-assembly process. We suggest that the Ri-boswitch-VLPs intervene earlier in the accumulation process by preventing some of the adenine to enter the yeast and then leading to fewer aggregates. To provide further support for this hypothesis, we used the ProteoStat dye, which stains amyloid fibrils and aggregates, and adenine intracellular aggregates to detect the presence of amyloid-like aggregates in the yeast model. The staining was followed by flow cytometry analysis, which allowed the detection of a decrease in the aggregation in the presence of the Ri-boswitch-VLPs (Figure 4e). Confocal microscopy further allowed the identification of the intracellular localization of the aggregates, showing stained dots only in the yeast model following the addition of adenine (Figure 4f). The stained dots were not detected in the presence of the Ri-boswitch-VLPs.

**Figure 4.**
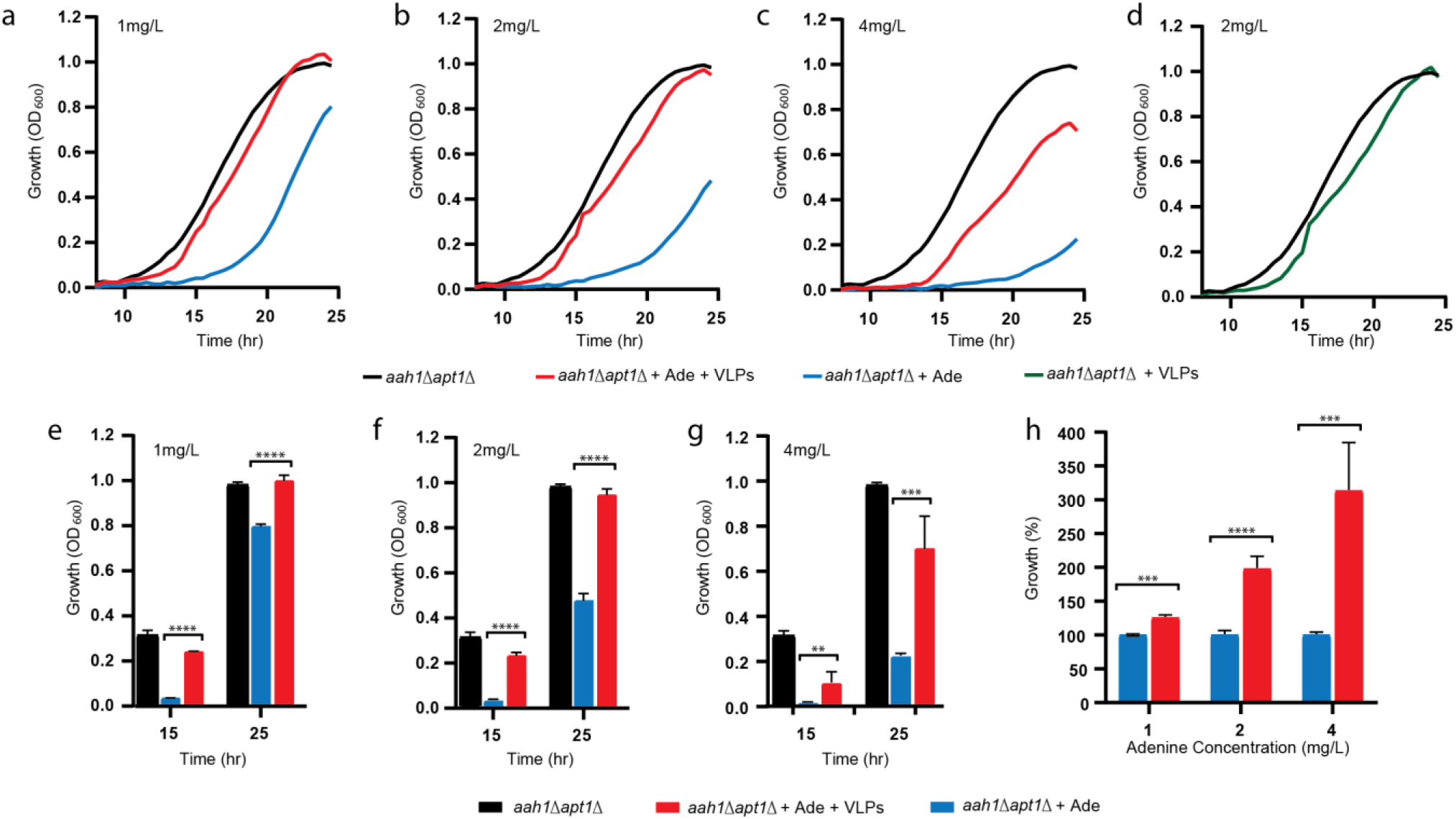
Riboswitch-VLPs supplementation to the adenine salvage in vivo yeast model. a-c. The aah1Δ apt1Δ mutant was grown without adenine (aah1Δ apt1Δ) or in the presence of (a) 1, (b) 2, and (c) 4 mg/L adenine (aah1Δ apt1Δ +Ade). Where indicated, the medium was supplemented with Riboswitch-VLPs (aah1Δ apt1Δ + Ade + VLPs). d. The aah1Δ apt1Δ mutant was grown without adenine and with Riboswitch-VLPs (aah1Δ apt1Δ + VLPs). e-g. Growth of the aahlΔ aptlΔ mutant without adenine (aah1Δ apt1Δ) or in the presence of (e) 1, (f) 2, and (g) 4 mg/L adenine (aah1Δ apt1Δ +Ade) at different time points. Where indicated, the medium was supplemented with Riboswitch-VLPs (aah1Δ apt1Δ + Ade + VLPs). h. Percentage of growth of the mutant yeast with adenine supplemented with Riboswitch-VLPs (aah1Δ apt1Δ + Ade + VLPs) compared to the nonsupplemented mutant (aah1Δ apt1Δ + Ade) at T=25 hours.

**Figure 5.**
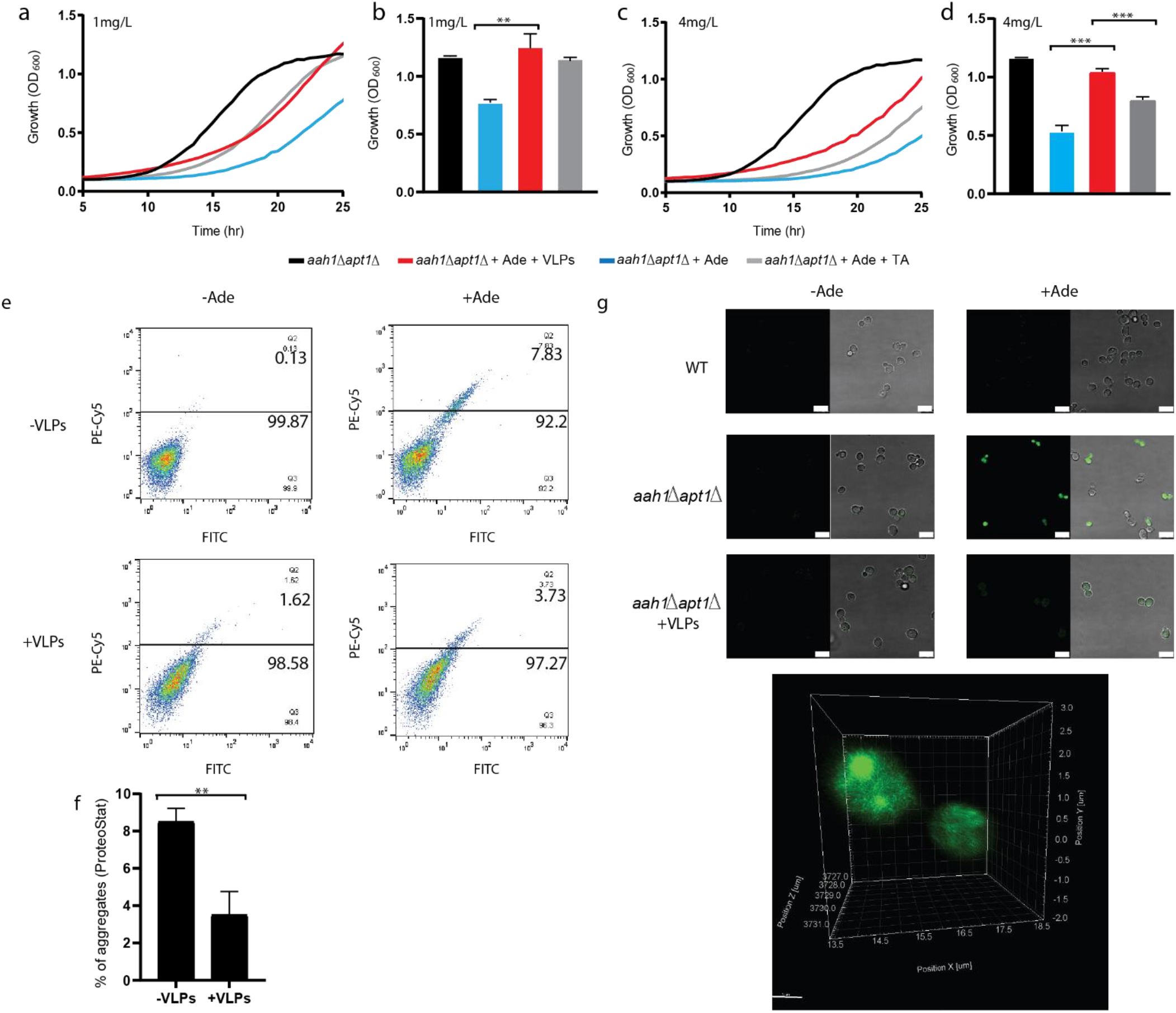
Comparison between Riboswitch-VLPs and TA on in vivo aggregation. a-d. The aahlΔ aptlΔ mutant was grown without adenine or in the presence of (a, b) 1 mg/L and (c, d) 4 mg/L adenine (aah1Δ apt1Δ +Ade). The medium was supplemented with either Riboswitch-VLPs (aah1Δ apt1Δ + Ade + VLPs) or 0.5mM TA (aah1Δ apt1Δ + Ade + TA). a, c. Growth curves. b, d. Endpoint graphs. e, f. Flow cytometry analysis of aah1Δ apt1Δ cells under the indicated conditions following ProteoStat staining. Adenine concentration was 2mg/L. **P < 0.005 (Student’s t-test). Values are the mean ± s.d. of three experiments. g. Representative confocal and differential interference contrast (DIC) images of WT, aah1Δ apt1Δ and aah1 Δapt1Δ + VLPs cells following ProteoStat staining (green). Adenine concentration was 2mg/L. Scale bar is 7.5 μm and a Z-stack followed by 3D reconstruction of aah1Δ apt1Δ under the indicated conditions following ProteoStat staining.

In the past few years, it became clear that metabolite amyloids play a key role in inherited metabolic disorders, leading the way towards the development of new possible treatments such as the suggested Riboswitch NCs. In the described technology, the VLPs’ shell protects the functional RNA under cellular conditions and in the presence of added RNase and DNase. As a result, the encapsulated RNA does not require additional structural design or chemical modifications to increase its stability *in vivo*. The VLPs offer a convenient and effective way to package and protect different cargos that can also be delivered to a specific cell or tissue. These features are much needed for practical medical applications for metabolic disorders as well as other diseases.

## METHODS

### Plasmid preparation

A DNA sequence encoding the maturation protein and the coat protein was synthesized (IDT) and transformed into pACYC-DUET E. coli using the Golden Gate standard procedure. The non-coding RNA sequence was inserted using reverse PCR (primers were purchased from IDT).

### Induction and expression of Riboswitch-VLPs

The plasmid was transformed into E.coli BL21 (DE3) cells (Novagene), according to the manufacturer’s instructions. Bacterial cells were grown in LB broth Miller (Difco) containing chloramphenicol (25 μg/mL) (Gold Biotechnology, USA) at 37 °C to OD600 = 1.7. 10 mL of the bacterial culture were diluted 1:1000in 1L Terrific broth (TB, BactlLAB) containing 25μg/mL chloramphenicol and cultivated at 37°C to OD600 = 0.4-0.8. Protein expression was induced by the addition of 1mM isopro-pyl-L-thio-D-galactopyranoside (IPTG) (Gold Biotechnology) at 37°c and after 35 minutes, rifampicin (Gold Biotechnology) was added to a final concentration of 200 μg/mL. The culture was further cultivated at 37°C for 16 hours. The cell suspension was centrifuged at 4000g for 20 minutes at 4 °C and cells were washed once in 200 mL PBS buffer (Hylabs) and centrifuged at 4000g for 20 minutes at 4 °C. The pellet was resuspended in 4 mL sonication buffer (100 mM NaH2PO4, 600 mM NaCl, pH 8.0 (both from Sigma-Aldrich), complete EDTA-free protease inhibitor (Roche), 200U of DNase I, and 200 μL of 10 mg/mL RNase A (both from Thermo Fisher Scientific)). The cells were then lysed using ultrasonic disruption for a total of 120 seconds, 20% amplitude (Disintegrator Sonicator W385). To eliminate cell debris, the lysed bacterial suspension was centrifuged at 12000 RPM for 15–30 minutes at room temperature. The supernatant containing Riboswitch-VLPs was filtered through a 0.22-μm syringe filter (Merck).

### Purification of Riboswitch-VLPs using Ni bead columns

The filtered supernatant was mixed at a 1:1 ratio with 2× binding buffer (100 mM NaH2PO4, 600 mM NaCl, 30 mM imidazole, pH=8.0, all from Sigma-Aldrich). Subsequently, His-tagged Ri-boswitch-VLPs were purified using a Ni beads gravity column loaded with high-performance Ni beads (GE Healthcare). Briefly, the column was pre-equilibrated with 50 mL binding buffer (50 mM NaH2PO4, 300 mM NaCl, 15 mM imidazole, pH 8.0, all from Sigma-Aldrich). The His-tagged Riboswitch-VLPs bound to the column were subsequently washed with 100 mL washing buffer (50 mM NaH2PO4, 300 mM NaCl, 30 mM Imidazole) and eluted from the column using10 mL elution buffer (50 mM NaH2PO4, 300 mM NaCl, 200 mM imidazole, pH 8.0, all from Sigma-Aldrich). All the eluted fractions were tested using a Bradford reagent assay (diluted 1:5 with ddH2O) (190 μL Bradford, 10 μL sample) (Bio-rad). Positive fractions were desalted using 100 kDa amicons (Merck) and converted first to ultrapure water and then to riboswitch stabilizing buffer (10 mM MgCl2 (Merck), 50 mM Tris-HCl (Bio-Lab) (pH 8.3), and 100 mM KCl (Merck)) for 24 hours each.

### *In vitro* transcription

A 1000 nucleotide transcript was transcribed using the Ribo-probe Combination System--SP6/T7 RN (Promega), according to the manufactures instructions. The transcript was then purified using the MEGAclear™ Transcription Clean-Up Kit (Thermo Fisher Scientific).

### Nuclease resistance assay

Riboswitch-VLPs, as well as in vitro transcribed naked RNA, were incubated with and without 10U of RNase A (Thermo Fisher Scientific) at 37 °C for 1 hour. Following incubation, the samples were analyzed using 1% agarose gel electrophoresis.

### Yeast growth assays

The strains used in this work were BY4741 (WT) and the adenine salvage mutant, aah1Δ apt1Δ, that was recently established by a disruption of both AAH1 and APT1.18 Yeast strains were grown overnight at 30 °C in a synthetic defined SD medium (6.7g/L Yeast Nitrogen Base without Amino Acids (Difco) and 20g/L glucose (Merck)) containing a specific mixture of amino acids and nucleobases. Strains were cultured in SD medium with adenine (adenine hemisulfate) at different concentrations as indicated (+ Ade) or without adenine (-Ade) as well as with Riboswitch-VLPs (+ Ade + VLPs).

For OD600 measurements, cells were diluted to OD600=0.01. 200 μL of cells were plated on 96 wells microplates (Greiner, F-BOTTOM) with increasing concentrations of adenine (1, 2 and 4 mg/L) and with 2.5% (V/V) of the Riboswitch-VLPs and incubated at 30 °C for 25 h with continuous shaking. OD600 was measured using TecanTM SPARK 10M plate reader. The results displayed are representative of three biological experiments performed in triplicates.

### TEM

TEM imaging was performed by applying 10μL samples onto 400-mesh copper grids covered by a carbon-stabilized Formvar film (SPI, West Chester, PA, USA). The samples were allowed to adsorb for 2 min before excess fluid was blotted off. Negative staining was then achieved by depositing 10μL of 2% uranyl acetate on the grid for 2 min before blotting off excess fluid. Micrographs were recorded using a Tecnai 12 electron microscope (FEI, Tokyo, Japan) operating at 120 kV.

### DLS

For DLS measurements, 1 mL of purified Riboswitch-VLPs were used. Particles size was measured using Malvern Panalytical Zetasizer Nano ZS (Light Source: He-Ne laser 633nm, Max 5mW) with a protocol specified for protein in water solution. Measurements of a sample were performed in triplicates in succession, and the reported values are the calculated Z average diameter and peak 1 average with standard deviation as error bars.

### Flow cytometry

Following incubation with adenine and VLPs (as outlined above) 1 mL of logarithmic cells were washed with PBS buffer and sonicated using 15 s pulses at 20% power. For each sample, 2*106 cells were resuspended with ProteoStat dye that was prepared according to the manufacturer’s instructions (Enzo Life Sciences) and diluted 1:3000 in PBS. Cells were incubated for 15 min at room temperature protected from light. Flow cytometry was performed using Stratedigm S1000EXi and the CellCapTure software (Stratedigm, San Jose, CA). Live cells were gated (P1) by forward scatter and side scatter. Fluorescence channels for FITC (530/30) and PE-Cy5 (676/29) were used utilizing a 488 nm laser source. A total of 50,000 events were acquired for each sample. Analyses were performed using the FlowJo software (TreeStar, version 10). The results displayed are representative of three biological experiments performed in triplicates.

### Confocal images

One milliliter of logarithmic cells incubated with adenine and VLPs (as mentioned above) was washed with PBS buffer and sonicated using 15 s pulses at 20% power and resuspended in 50 μL of ProtesoStat dye diluted 1:250 in PBS. Cells were incubated for 15 min at room temperature protected from light. 10 μL of each sample was deposited on poly-lysine-coated glass slides (Sigma-Aldrich). Cells were imaged using Leica TCS SP8 laser confocal microscope with ×63 1.4 NA or ×100 1.4 NA oil objectives. An argon laser with 488 excitation line was used for ProteoStat (emission wavelength, 500–600 nm). The results displayed are representative of three biological experiments

## ASSOCIATED CONTENT

### Supporting Information

TR-RNA and riboswitch structrers, TEM images of WT MS2 and VLPs encapsulating random RNA, supplementation of Riboswitch-VLPs at different growth, flow cytometry analysis of WT yeast and model yeast supplementation of VLPs with and without adenine, plasmid maps, DNA sequances’ table. (Figures S1-S8), Table S1. This material is available free of charge via the Internet at http://pubs.acs.org

## AUTHOR INFORMATION

### Author Contributions

S.Z.T. and D.G. designed and performed the experiments. H.A. and D.L.B.Y. assisted in designing experiments. M.G. and R.B. helped with the flow cytometry experiments. D.G. and D.L.B.Y. edited the manuscript. S.Z.T. and E.G. conceived the project, analyzed data and wrote the manuscript.

## ACKNOWLEDGMENT

S.Z.T was supported by the Marian Gertner Institute for Medical Nanosystems. The authors would like to thank Dr. S. Lichtenstein for the help with confocal microscopy, Dr. O. Sagi-Assif for help with the FACS analysis, D. Greitzer for graphical assistance, and members of the Gazit laboratory for helpful discussions.

## ABBREVIATIONS

WT: wild type
TR-RNA: translational repression RNA
VLPs: virus like particles
APRT: adenine phosphoribosyltransferase
ADA: adenosine deaminase
IEM: inborn error of metabolism
TEM: transmission electron microscopy
NCs: nanocarriers
TA: Tannic Acid

