## Supplemental information for "Engineered Riboswitch Nano-carriers as a Possible Disease-Modifying Treatment for Metabolic Disorders"

[1]The Shmunis School of Biomedicine and Cancer Research, Tel Aviv University, Tel Aviv 6997801, Israel

[2]BLAVATNIK CENTER for Drug Discovery, Tel Aviv University, Tel Aviv 6997801, Israel

[3]Department of Materials Science and Engineering Iby and Aladar Fleischman Faculty of Engineering, Tel Aviv University, Tel Aviv 6997801, Israel

DNA sequences' table ........................................................................................................................ **11-12**

References .................................................................................................................................................. **13**

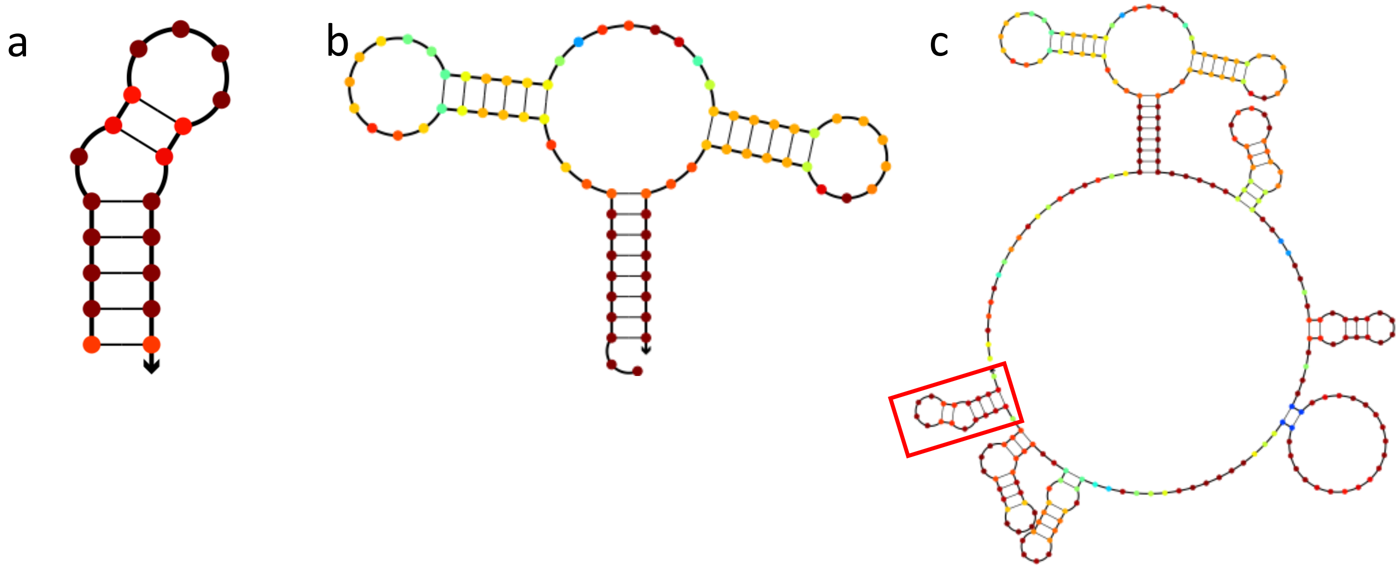

**Figure S1.** RNA secondary structures calculated by NUPACK [1, 2] of: a. TR RNA activator, b. Adenine Riboswitch A58U, and c. Adenine Riboswitch A58U conjugated to the TR RNA activator. (Red box indicates the TR RNA part.)

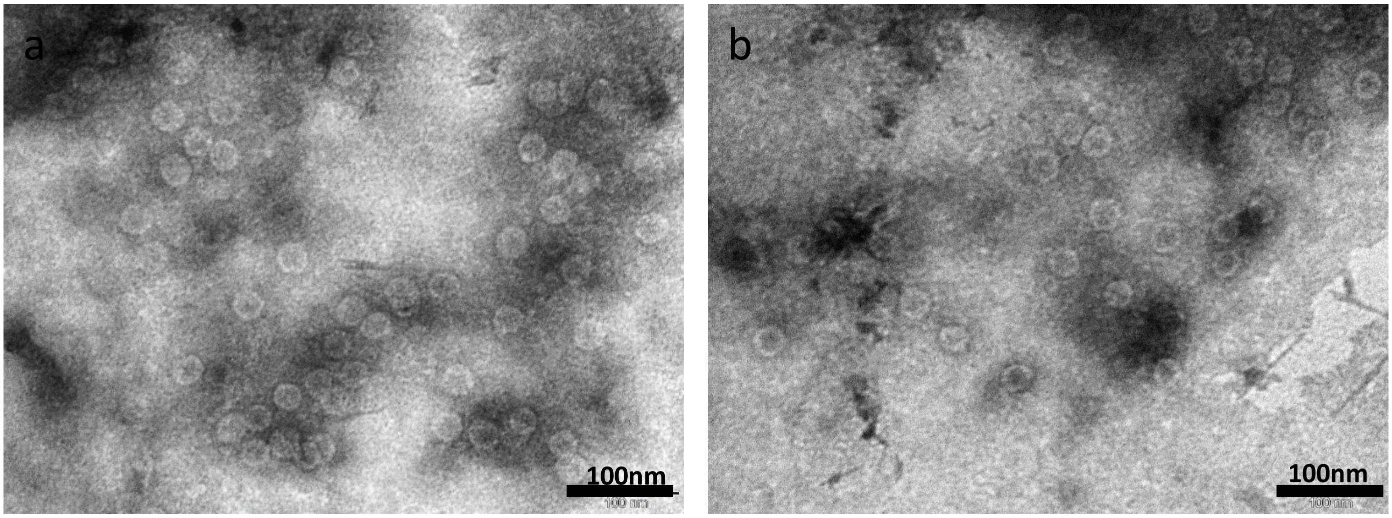

**Figure S2.** a and b. Transmission electron micrographs (TEM) of WT MS2.

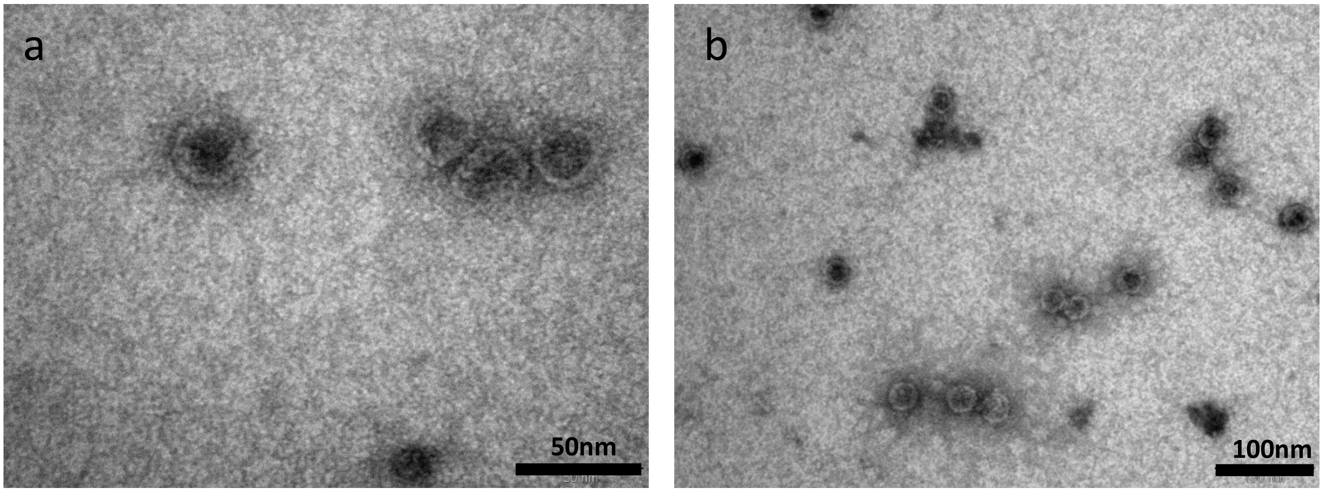

**Figure S3.** a and b. TEM images of VLPs-RNA nano-carriers encapsulating random RNA sequence containing the TR-RNA activator sequence.

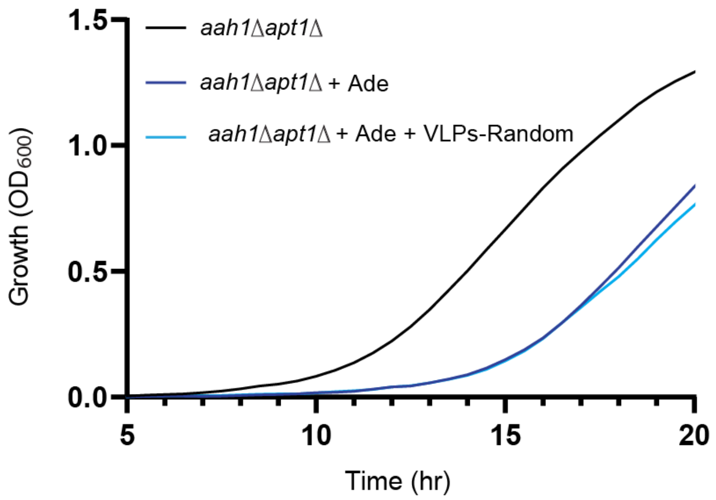

**Figure S4.** Growth curves of the yeast model (denoted as *aah1*Δ *apt1*Δ) with adenine (*aah1*Δ *apt1*Δ + Ade) and with VLPs-RNA nano-carriers encapsulating random RNA sequence (*aah1*Δ *apt1*Δ +Ade + VLPs-Random).

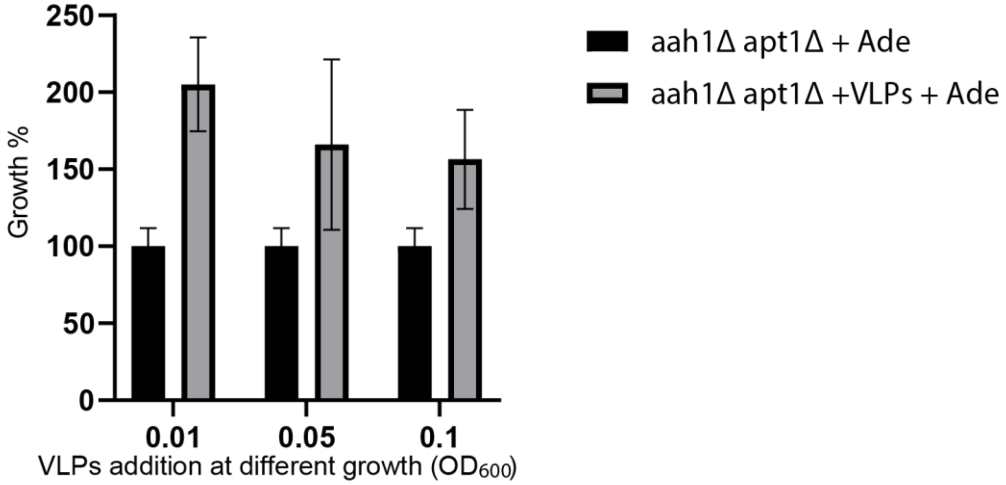

**Figure S5.** Addition of the VLPs-Riboswitch at different $OD_{600}$ values of the model yeast model in the presence of adenine.

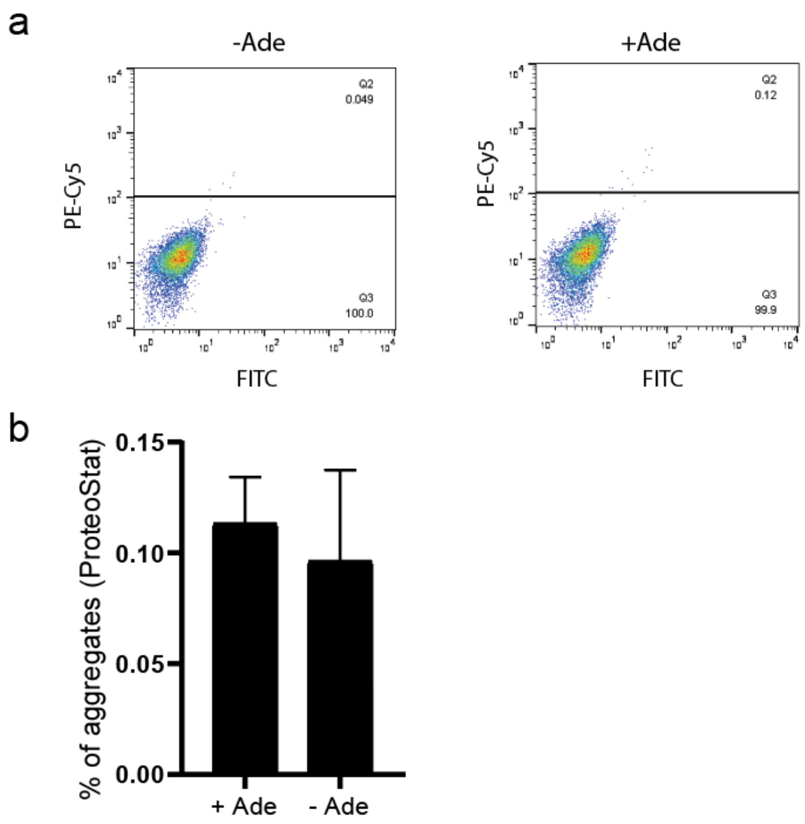

**Figure S6.** Flow cytometry analysis of WT cells under the indicated conditions following ProteoStat staining.

a

b

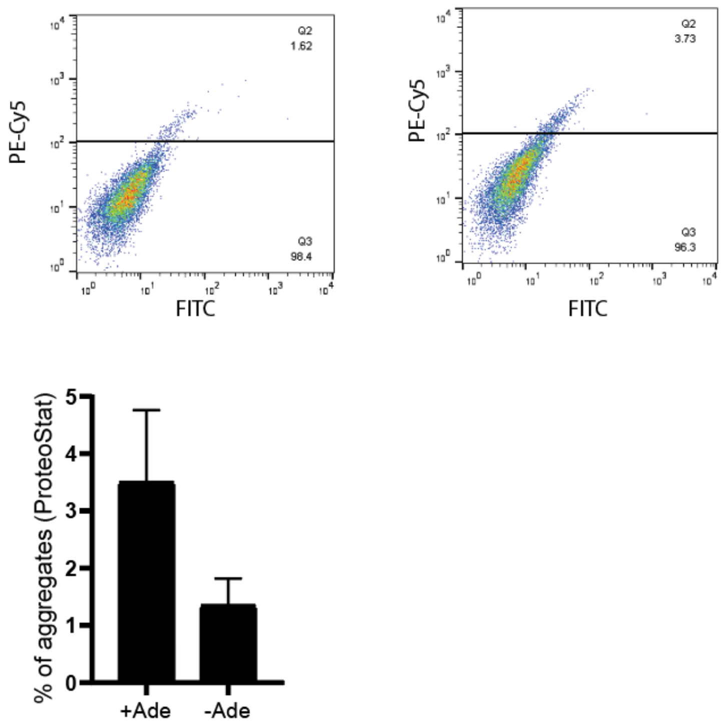

**Figure S7.** Flow cytometry analysis of *aah1*Δ *apt1*Δ cells with VLPs-Riboswitch under the indicated conditions following ProteoStat staining.

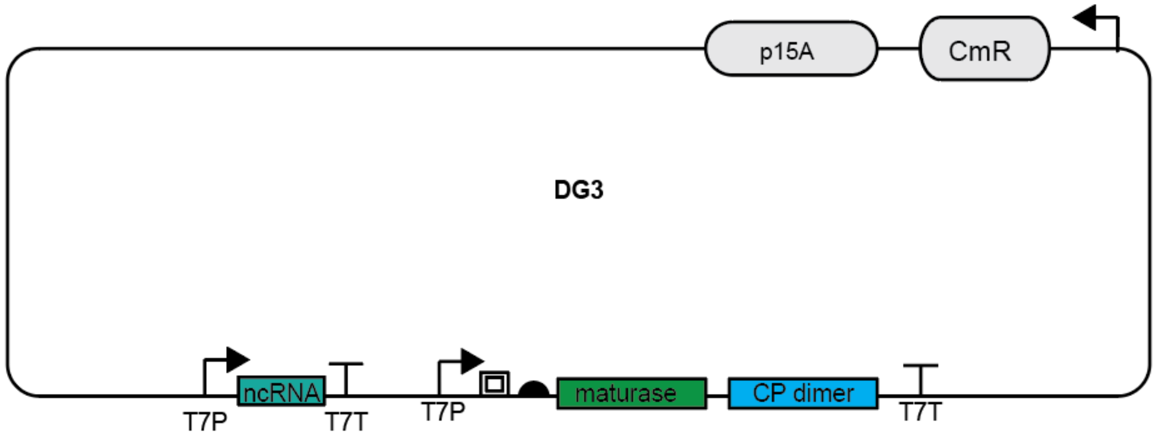

**Figure S8.** Plasmid map of the co-expression of the MS2 CP and the ncRNA.

**Table S1.** DNA sequences.

| Part | Sequence |
| --- | --- |
| TR RNA | ACATGAGGATCACCCATGT |
| TMpac | CGCTTCCGTCAAACCCCTGCGCGTTGTATAACCTCAATAATATGGTTTGAG<br>GGTGTCTACCAGGAACCGTAAATTCCTGATTACAACGCAAACCGGATGAT<br>AGACCTCACCTCCCCGCCCAATACTGAAATCTCATTAATACGCATACCCC<br>CACTATACACACGCAATCACCACATTAGCACAATGAATAATCATCGTACG<br>GGAGAAAACATTCTAAACCCACATGAGGATCACCCATGT |
| Adenine riboswitch (A58U) | GCGCGTTGTATAACCTCAATAATATGGTTTGAGGGTGTCTACCAGGAACC<br>GTAAATTCCTGATTACAACGC |
| Maturase and CP dimer gblock | ACGACTCACTATAGGGGAATTGTGAGCGGATAACAATTCCCCTGTAGAAA<br>TAATTTTGTTTAACTTTAATAAGGAGATATACCATGGTGGCTATCGCTGTA<br>GGTAGCCGGAATTCCATTCCTAGGAGGTTTGACCTGTGCGAGCTTTTAGT<br>ACCCTTGATAGGGAGAACGAGACCTTCGTCCCCTCCGTTCGCGTTTACGC<br>GGACGGTGAGACTGAAGATAACTCATTCTCTTTAAAATATCGTTCGAACT<br>GGACTCCCGGTCGTTTTAACTCGACTGGGGCCAAAACGAAACAGTGGCAC<br>TACCCCTCTCCGTATTCACGGGGGGCGTTAAGTGTCACATCGATAGATCA<br>AGGTGCCTACAAGCGAAGTGGGTCATCGTGGGGTCGCCCGTACGAGGAG<br>AAAGCCGGTTTCGGCTTCTCCCTCGACGCACGCTCCTGCTACAGCCTCTTC<br>CCTGTAAGCCAGAACTTGACTTACATCGAAGTGCCGCAGAACGTTGCGAA<br>CCGGGCGTCGACCGAAGTCCTGCAAAAGGTCACCCAGGGTAATTTTAACC<br>TTGGTGTTGCTTTAGCAGAGGCCAGGTCGACAGCCTCACAACTCGCGACG<br>CAAACCATTGCGCTCGTGAAGGCGTACACTGCCGCTCGTCGCGGTAATTG<br>GCGCCAGGCGCTCCGCTACCTTGCCCTAAACGAAGATCGAAAGTTTCGAT<br>CAAAACACGTGGCCGGCAGGTGGTTGGAGTTGCAGTTCGGTTGGTTACCA<br>CTAATGAGTGATATCCAGGGTGCCTATGAGATGCTTACGAAGGTTCACCT<br>TCAAGAGTTTCTTCCTATGAGAGCCGTACGTCAGGTCGGTACTAACATCA<br>AGTTAAATGGCCGTCTGTCGTATCCAGCTGCAAACTTCCAGACAACGTGC<br>AACATATCGCGACGTATCGTGATATGGTTTTACATAAACGATGCACGTTT<br>GGCATGGTTGTCGTCTCTAGGTATCTTGAACCCACTAGGTATAGTGTGGG<br>AAAAGGTGCCTTTCTCATTCGTTGTCGACTGGCTCCTACCTGTAGGTAACA<br>TGCTCGAGGGCCTTACGGCCCCCGTGGGATGCTCCTACATGTCAGGAACA<br>GTTACTGACGTAATAACGGGTGAGTCCATCATAAGCGTTGACGCTCCCTA<br>CGGGTGGACTGTGGAGAGACAGGGCACTGCTAAGGCCCAAATCTCAGCC<br>ATGCATCGAGGGGTACAATCCGTATGGCCAACAACTGGCGCGTACGTAAA<br>GTCTCCTTTCTCGATGGTCCATACCTTAGATGCGTTAGCATTAATCAGGCA<br>ACGGCTCTCTAGATAGAGCCCTCAACCGGAGTTTGAAGCATGGCTTCTAA<br>CTTTACTCAGTTCGTTCTCGTCGACAATGGCGGAACTGGCGACGTGACTGT<br>CGCCCCAAGCAACTTCGCTAACGGGGTCGCTGAATGGATCAGCTCTAACT<br>CGCGTTCACAGGCTTACAAAGTAACCTGTAGCGTTCGTCAGAGCTCTGCG<br>CAGAATCGCAAATACACCATCAAAGTCGAGGTGCCTAAAGTGGCAACCC<br>AGACTGTTGGTGGTGTAGAGCTTCCTGTAGCCGCATGGCGTTCGTACTTA<br>AATATGGAACTAACCATTCCAATTTTCGCTACGAATTCCGACTGCGAGCTT<br>ATTGTTAAGGCAATGCAAGGTCTCCTAAAAGATGGAAACCCGATTCCCTC<br>AGCAATCGCAGCAAACTCCGGCATCTACGCTAACTT |
| CP dimer | AGCCCTCAACCGGAGTTTGAAGCATGGCTTCTAACTTTACTCAGTTCGTTC<br>TCGTCGACAATGGCGGAACTGGCGACGTGACTGTCGCCCCAAGCAACTTC<br>GCTAACGGGGTCGCTGAATGGATCAGCTCTAACTCGCGTTCACAGGCTTA<br>CAAAGTAACCTGTAGCGTTCGTCAGAGCTCTGCGCAGAATCGCAAATACA<br>CCATCAAAGTCGAGGTGCCTAAAGTGGCAACCCAGACTGTTGGTGGTGTA<br>GAGCTTCCTGTAGCCGCATGGCGTTCGTACTTAAATATGGAACTAACCAT<br>TCCAATTTTCGCTACGAATTCCGACTGCGAGCTTATTGTTAAGGCAATGCA<br>AGGTCTCCTAAAAGATGGAAACCCGATTCCCTCAGCAATCGCAGCAAACT<br>CCGGCATCTACGCTAACTTTACTCAGTTCGTTCTCGTCGACAATGGCGGTA<br>CCCATCACCATCACCATCACGGTACCGGCGACGTGACTGTCGCCCCAAGC |

AACTTCGCTAACGGGGTCGCTGAATGGATCAGCTCTAACTCGCGTTCACA
GGCTTACAAAGTAACCTGTAGCGTTCGTCAGAGCTCTGCGCAGAATCGCA
AATACACCATCAAAGTCGAGGTGCCTAAAGTGGCAACCCAGACTGTTGGT
GGTGTAGAGCTTCCTGTAGCCGCATGGCGTTCGTACTTAAATATGGAACT
AACCATTCCAATTTTCGCTACGAATTCCGACTGCGAGCTTATTGTTAAGGC
AATGCAAGGTCTCCTAAAAGATGGAAACCCGATTCCCTCAGCAATCGCAG
CAAACTCCGGCATCTACTAATAGACGCCGG

1. Dirks, R. M.; Pierce, N. A., A partition function algorithm for nucleic acid secondary structure including pseudoknots. *Journal of computational chemistry* **2003,** *24* (13), 1664-77.
2. Dirks, R. M.; Pierce, N. A., An algorithm for computing nucleic acid base-pairing probabilities including pseudoknots. *Journal of computational chemistry* **2004,** *25* (10), 1295-304.
